# Phosphorylation of MYCN and MAX by PAK family kinases is a novel tumor-suppressor mechanism in neuroblastoma with potential therapeutic implications

**DOI:** 10.1101/2023.12.04.570005

**Authors:** Esteban J Rozen, Kim Wigglesworth, Jason Shohet

## Abstract

High-risk Neuroblastoma (HR-NB) is a very aggressive pediatric cancer, responsible of over 15% of all childhood cancer-associated deaths. Despite very aggressive multimodal interventions, less than 50% of HR-NB patients survive, exhibiting serious long-term sequelae from therapy. Therefore, more efficient, and less toxic interventions are urgently needed. Genomic amplification of the MYCN gene is observed in about 50% of all HR-NB cases, and is the most reliable genomic hallmark associated to a bad prognosis. c-MYC is highly expressed in a significant amount of the remaining (non-MYCN-amplified) HR-NB cases. Here, we investigated an endogenous mechanism mediated by the PAK2 kinase and known to phosphorylate and suppress c-MYC transcriptional activity. We uncovered that PAK2 can also phosphorylate MYCN and its obligate transcriptional partner MAX in two conserved Ser/Thr residues, disrupting MYCN:MAX interaction, transcription, and neuroblastoma cell proliferation. We further provide evidence for a potential mechanism by which PAK kinase activity is blunted in MYCN-amplified neuroblastoma tumors and propose an innovative strategy to circumvent such signaling impairment and potentially suppress HR-NB tumor growth.

## Introduction

Neuroblastoma is the most common extracranial cancer of the childhood, accounting for ∼15% of pediatric cancer mortality. High-risk neuroblastoma cases have a dramatically poor prognosis, with less than 50% of survival rates^1^. Genomic amplification of MYCN occurs in ∼50% of high-risk neuroblastomas (HR-NB) (Brodeur) and greatly contributes to therapy resistance and disease-related death^2^. Therefore, MCYN is considered the most consistent genomic hallmark of aggressive neuroblastoma^3–5^. Lack of effective treatments is still a major clinical challenge in the treatment of MYCN-amplified neuroblastoma. MYCN belongs to the MYC family of oncogenes, which includes three closely related genes, c-MYC, MYCN and MYCL^6,7^. Amplification of the MYCN gene is predominantly found in neuroendocrine tumors, including neuroblastoma, medulloblastoma, retinoblastoma and neuroendocrine prostate cancer^8–10^. MYCN amplification is an initiating event, driving the development of high-risk neuroblastoma. Targeted MYCN overexpression in peripheral neural crest is sufficient to initiate disease in mouse models^11^, while MYCN inhibition broadly reverses tumor stem–like phenotypes and aberrant proliferation^12^. Hence, therapeutic strategies that target MYCN expression or protein stability reduce tumor growth in preclinical tumor models^13–15^. Alternatively, a significant amount of MYCN-non-Amplified HR cases (possibly up to 25%) express relatively high levels of c-MYC^16,17^ in a mutually exclusive fashion^17^. The MYC family of oncoproteins are also referred to as ‘super-transcription factors’, as they can induce broad transcriptional responses associated with virtually every aspect of cancer biology. Both c-MYC and MYCN have been found to largely remodel the cancer cis-regulatory landscape, leading to global transcriptional amplification^18,19^.

## Results

The MYC family of oncogenes belong to the basic helix-loop-helix leucine zipper (bHLH/Z) transcription factor superfamily, and need to form heterodimers with their binding partner MAX for DNA recognition and transcriptional modulation^20–22^. Binding of MYC-MAX dimers to DNA occurs at specific motifs -CACGTG-known as E-boxes^22,23^, after which accessory transactivators are recruited^24^. c-MYC transcriptional activity is tightly regulated at many levels, particularly, at the post-transcriptional, by means of phosphorylations that suppress its interaction with other proteins or the DNA, or that induce its degradation^25–27^. Among these, the evolutionarily conserved residues Thr-358, Ser-373, and Thr-400 have been characterized, as phosphorylation targets of the stress-induced kinase PAK2^28,29^. Thr-358 is located in the basic region of the bHLH domain, which interacts directly with DNA, whereas Ser-373 and Thr-400 are located in the adjacent helix– loop–helix region, which participates in the interaction with MAX (Fig. 1A). Phosphorylation of these residues by PAK2 has been shown to interfere with formation of the c-MYC–MAX–DNA ternary complex and promote an E-box-independent transcription and differentiation^28–30^. In this context, Aznar-Benitah *et al*.^31^ suggested a Rac1/PAK2-dependent inhibitory phosphorylation of c-MYC as a major modulator of epidermal stem cell differentiation. Later, Uribesalgo *et al*.^30^ demonstrated a role for PAK2-mediated c-MYC phosphorylation in the functional switch of this transcription factor from pro-leukemogenic (through its interaction with MAX) to a tumor-suppressive activity (through a MAX-independent interaction with Retinoic Acid Receptor α). Importantly, the unrelated protein kinase PKCζ was also shown to phosphorylate c-MYC Ser373, again suppressing c-MYC oncogenic activity in prostate cancer models^32^.

**Figure 1.**
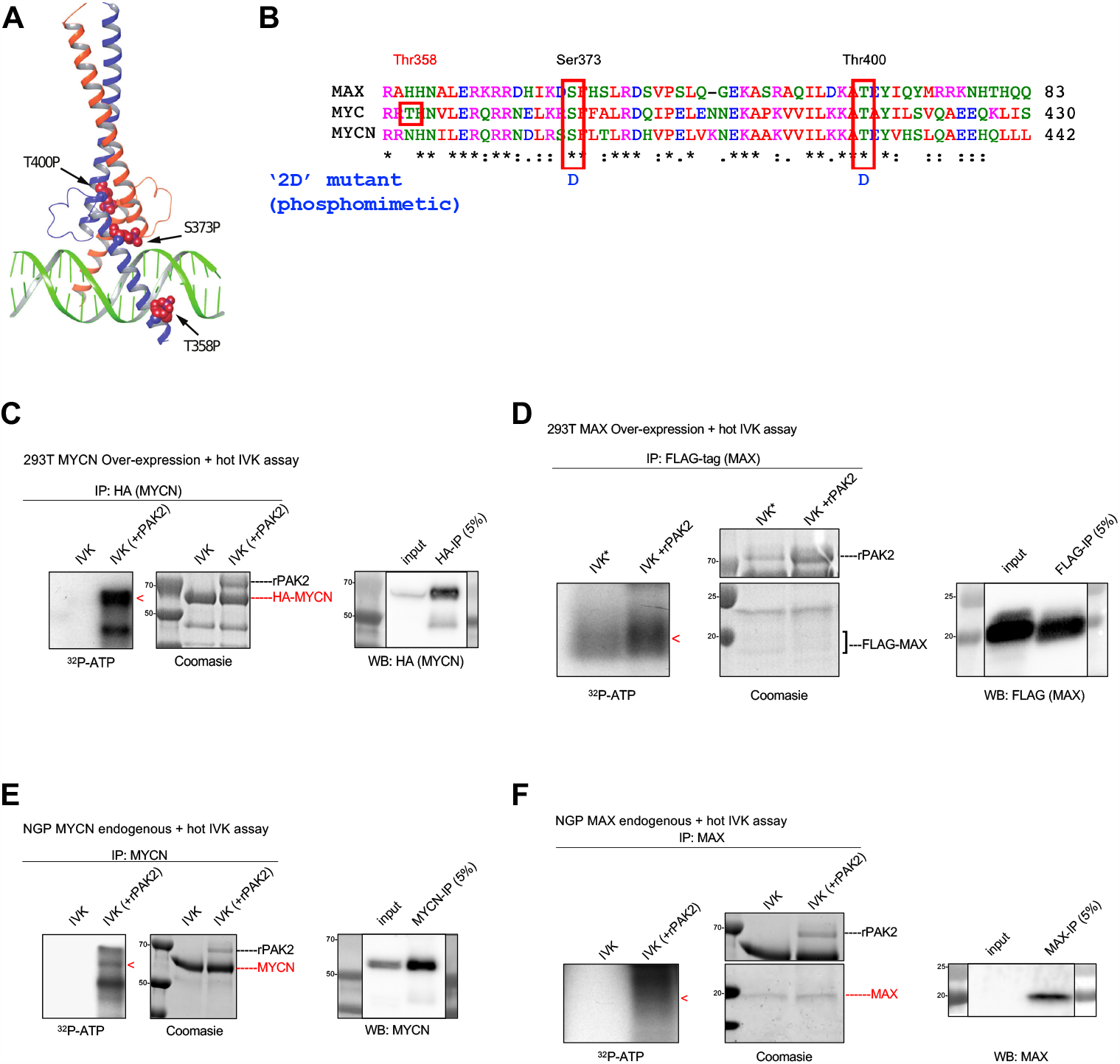
**A**. Reproduced without modifications from Figure 1A of Macek *et al*. (2108)^29^: Crystal structure of the heterodimer formed by the bHLH/Z domains of Myc (blue) and Max (orange) in complex with E-box DNA (PDB code 1NKP), showing potential phosphorylation sites as red space-filling spheres. **B**. Amino-acid sequence alignment of human c-MYC 356-430 and the corresponding homologous sequences of human MAX (25-83) and human MYCN (356-442), depicting the conservation of PAK2 target sites (red boxes) and the corresponding mutations to Aspartic Acid (D) in the ‘2D’ mutant versions. **C**. Cell lysates from HEK 293T cells overexpressing HA-MYCN WT were subjected to HA-immuno-precipitation. HA-immuno-complexes were then subjected to *in vitro* kinase (IVK) assay with radioactive [γ-^32^P]-ATP in the absence or presence of constitutively activated mutant recombinant PAK2 (T402E) protein (rPAK2). The left panel shows the autoradiographic signal (^32^P-ATP) for phosphorylated HA-MYCN (red arrowhead). The middle panel shows the corresponding Coomassie staining imaging of the same gel, showing the bands corresponding to HA-MYCN and rPAK2. The right panel shows Western blot analysis with anti-HA (MYCN) for the input sample and a small aliquot (5%) of the HA-immuno-precipitate. **D**. Cell lysates from HEK 293T cells overexpressing FLAG-MAX WT were subjected to FLAG-immuno-precipitation. FLAG-immuno-complexes were then subjected to *in vitro* kinase (IVK) assay with radioactive [γ-^32^P]-ATP in the absence or presence of constitutively activated mutant recombinant PAK2 (T402E) protein (rPAK2). The left panel shows the autoradiographic signal (^32^P-ATP) for phosphorylated FLAG-MAX* (red arrowhead). The middle panel shows the corresponding Coomassie staining imaging of the same gel, showing the bands corresponding to FLAG-MAX and rPAK2*. The right panel shows Western blot analysis with anti-FLAG (MAX) for the input sample and a small aliquot (5%) of the FLAG-immuno-precipitate. **\*** The asterisk denotes that there was a slight leakage of sample from the rPAK2-conatining lane into the rPAK2-null lane, which can be observed by the presence of a fainter rPAK2 band in the Coomasie staining imaging and by the presence of a fainter but detectable ^32^P-ATP autoradiographic signal in the supposedly negative control. **E**. Cell lysates from NGP MYCN-positive neuroblastoma cells were subjected to MYCN-immuno-precipitation. MYCN-immuno-complexes were then subjected to *in vitro* kinase (IVK) assay with radioactive [γ-^32^P]-ATP in the absence or presence of constitutively activated mutant recombinant PAK2 (T402E) protein (rPAK2). The left panel shows the autoradiographic signal (^32^P-ATP) for phosphorylated MYCN (red arrowhead). The middle panel shows the corresponding Coomassie staining imaging of the same gel, showing the bands corresponding to MYCN and rPAK2. The right panel shows Western blot analysis with anti-MYCN for the input sample and a small aliquot (5%) of the MYCN-immuno-precipitate. **F**. NGP MYCN-positive neuroblastoma cells were subjected to MAX-immuno-precipitation. MAX-immuno-complexes were then subjected to *in vitro* kinase (IVK) assay with radioactive [γ-^32^P]-ATP in the absence or presence of constitutively activated mutant recombinant PAK2 (T402E) protein (rPAK2). The left panel shows the autoradiographic signal (^32^P-ATP) for phosphorylated MAX (red arrowhead). The middle panel shows the corresponding Coomassie staining imaging of the same gel, showing the bands corresponding to MAX and rPAK2. The right panel shows Western blot analysis with anti-MAX for the input sample and a small aliquot (5%) of the FLAG-immuno-precipitate.

MYCN and c-MYC show a high degree of sequence and functional conservation^33^, as demonstrated by Malynn *et al*.^34^, who generated mice in which the endogenous c-Myc coding sequences were replaced by N-Myc sequence. Strikingly, mice homozygous for this replacement mutation could survive into adulthood and reproduce. With this in mind, we wondered whether the residues phosphorylated by PAK2 on c-MYC - Thr-358, Ser-373, and Thr-400-could also be conserved in MYCN and possibly other members of the bHLH/Z family of transcription factors. Sequence alignment showed that Ser-373 and Thr-400 indeed are conserved in MYCN and MAX (Figure 1B), but not in other members of the bHLH/Z family (Supplementary figure 1). Based on this observation, we hypothesized that PAK2 might also phosphorylate MYCN and MAX, in the corresponding conserved residues. To this end, we overexpressed HA-MYCN and FLAG-MAX in HEK 293T cells. Whole cell lysates where then immuno-precipitated with either HA- or FLAG-bound magnetic beads, respectively, and immuno-precipitates were subjected to *in vitro* kinase (IVK) assays with radioactive [γ-^32^P]-ATP in the absence or presence of constitutively activated mutant recombinant PAK2 (T402E) protein. By this approach we could observe a clear autoradiographic signal corresponding to HA-MYCN (Figure 1C) and FLAG-MAX (Figure 1D) only in those samples in which exogenous recombinant PAK2 (rPAK2) was added, suggesting the direct phosphorylation MYCN and MAX by PAK2. To further confirm this observation, we next immuno-precipitated endogenous MYCN and MAX proteins from the MYCN-amplified neuroblastoma cell line NGP and performed rPAK2-mediated IVKs. Again, we could observe autoradiographic signals corresponding to endogenous MYCN (Figure 1E) and MAX (Figure 1F) only in the presence of activated PAK2, corroborating the hypothesis that MYCN and MAX can indeed be phosphorylated by PAK2 *in vitro*.

PAK2 phosphorylation of c-MYC on Ser373 and Thr400 has been shown to disrupt c-MYC:MAX interaction and subsequent transcriptional activity^28–30^. Therefore, we next hypothesized that PAK2 phosphorylation of MYCN and MAX on the correspondingly conserved residues (MYCN Ser400 and Thr427, and MAX Ser42 and Thr68), should also reduce their ability to interact with each other. To test this, we synthesized ‘2D’ mutant versions of both MYCN and MAX, in which the two mentioned Ser and Thr residues were mutated to aspartic acid (D), mimicking constitutively phosphorylated Ser/Thr residues (Figure 1B), as previously shown for c-MYC^28,30^. We then overexpressed the wild-type (WT) or mutated (2D) versions of the two proteins in different possible combinations and subjected them to immuno-precipitation assays against HA (MYCN; Figure 2A, left panel) or FLAG (MAX, Figure 2A, middle panel) and assessed the relative co-precipitation of the corresponding binding partner. As seen in Figure 2A (left panel), co-precipitation of MAX with MYCN immuno-complexes was maximal when both proteins were in their WT version, while significant reductions of co-precipitated MAX were seen when either MYCN or MAX had the 2D phospho-mimetic mutation. Interestingly, the interaction was minimal when both proteins carried the 2D mutations. Similar results were observed when MAX was immuno-precipitated and MYCN co-precipitation was analyzed (Figure 2A, middle panel), leading to the conclusion that phosphorylation of MYCN and MAX by PAK2 likely results in a significant reduction of MYCN:MAX interaction. Finally, we predicted that the observed reduction of MYCN:MAX interaction due to the PAK2 phospho-mimetic 2D mutations should also result in decreased transcriptional activity of the MYCN:MAX complex. To address this, we performed *in vitro* luciferase assays with the pEBOX4-luc reporter construct, harboring 4 tandem E-box elements controlling the expression of the luciferase reporter gene (pMyc4ElbLuc^35^). We transiently transfected HEK 293T cells with the reporter vector pEBOX4-luc along with and empty expression vector (‘pEBOX4-luc only’ control) or with the expression vectors encoding HA-MYCN and FLAG-MAX, either WT or 2D in all the possible combinations. As an additional control we transfected the pEBOX0-luc (no E-boxes) construct with the two WT versions of MYCN and MAX. As expected, the pEBOX0-luc and pEBOX4-luc only controls exhibited a minimal basal (∼10%) luciferase activity (Figure 2B). Concomitant expression of pEBOX4-luc with WT MYCN and WT MAX resulted in a ∼10-fold increase in luciferase activity (defined as 100%). Replacement of each WT version by its corresponding 2D mutant variant resulted in a significant reduction of the *in vitro* transcriptional activity, while simultaneous mutation of both factors resulted in nearly complete reduction of the luciferase activity, confirming that phosphorylation of MYCN and MAX in these conserved residues not only disrupts their interaction but also their transcriptional activity, as previously shown for c-MYC^28^. Interestingly, our co-IP and luciferase reporter results indicate an additive effect by the sequential phospho-mimetic mutation of MYCN and MAX, suggesting that the level of MYCN:MAX transcription inhibition is dependent on the level of PAK2 kinase activity.

**Figure 2.**
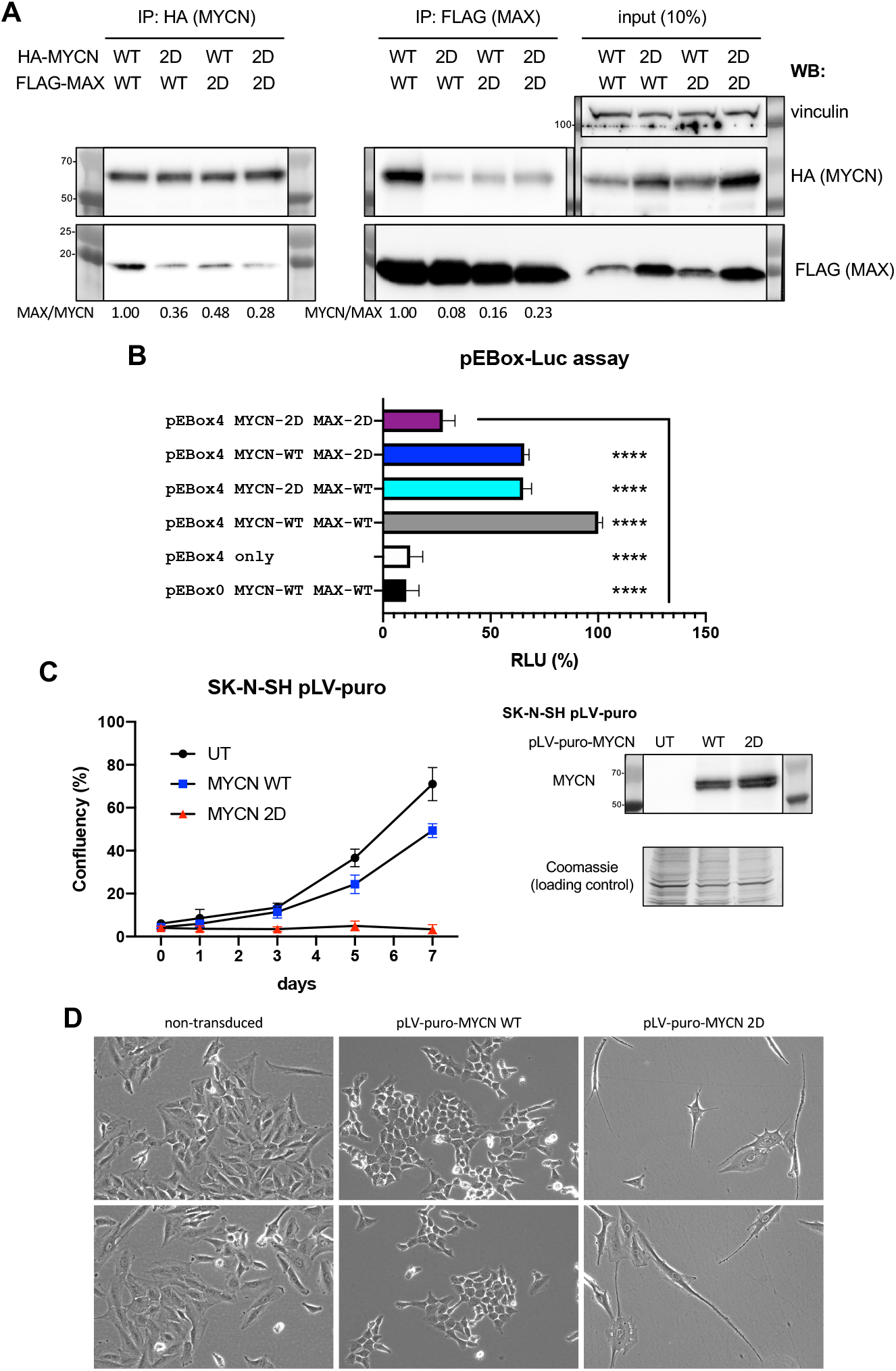
**A**. Left panel. Cell lysates from HEK 293T cells overexpressing HA-MYCN WT or 2D and FLAG-MAX WT or 2D in all 4 possible combinations were subjected to HA-immuno-precipitation. Immuno-complexes were resolved by SDS-PAGE and subjected to Western blot analysis for detection of HA-MYCN and the resulting co-precipitated FLAG-MAX. Middle panel. Cell lysates from HEK 293T cells overexpressing HA-MYCN WT or 2D and FLAG-MAX WT or 2D in all 4 possible combinations were subjected to FLAG-immuno-precipitation. Immuno-complexes were resolved by SDS-PAGE and subjected to Western blot analysis for detection of FLAG-MAX and the resulting co-precipitated HA-MYCN. Right panel. A small aliquot (10%) of the cell lysates from HEK 293T cells overexpressing HA-MYCN WT or 2D and FLAG-MAX WT or 2D in all 4 possible combinations were resolved by SDS-PAGE and subjected to Western blot analysis for detection of HA-MYCN, FLAG-MAX and the housekeeping gene product vinculin as a loading control. **B**. HEK 293T cells were transiently transfected with the reporter vector pEBOX4-luc along with and empty expression vector (‘pEBOX4-luc only’ control) or with the expression vectors encoding HA-MYCN and FLAG-MAX, either WT or 2D, in all the possible combinations. As an additional control we transfected the pEBOX0-luc (no E-boxes) construct with the two WT versions of MYCN and MAX. Luciferase activity is expressed in percentual Relative Luciferase Units (RLU %). Each condition was done in 5 technical replicates. **C**. Left panel. MYCN-negative SK-N-SH neuroblastoma cells were left untreated (UT) or transduced with lentiviral vectors for expression of the indicated MYCN version (WT vs. 2D). After puromycin selection, ∼2x10^3^ cells were plated on M96 well plates (4 replicates/condition) and allowed to grow for 7 days. Confluency of each well was measured every 1-2 days in an Image cytometer (Celigo, Nexcelom, see Materials and Methods section). Right panel. These same cells were also plated in parallel and processed for Western blot analysis to detect MYCN protein expression. Coomassie staining of the PVDF membrane was used as a global protein loading control after anti-MYCN staining. **D**. Representative images from C on day 5, demonstrating striking phenotypic changes in each condition.

Next, we wondered if the constitutive phosphorylation of MYCN would result in significant changes in the proliferative behavior of neuroblastoma cells. To this end, we engineered a lentiviral vector for constitutive overexpression of either MYCN WT or the 2D variant, under the puromycin selection gene (pLV-puro-MYCN WT/2D, Figure 2C right panel). We transduced the MYCN-negative neuroblastoma cell line SK-N-SH with these lentiviral particles, and upon selection, observed a striking reduction in the proliferative capacity of the MYCN 2D-expressing cells, which was not observed in those cells overexpressing MYCN WT or in the untransduced (UT) control cells (Figure 2C left panel). Microscopic analysis of these cells demonstrated a profound phenotypic change in the MYCN 2D cells, which exhibited enlarged and elongated cell bodies, reminiscent of cells undergoing senescence and/or neural differentiation (Figure 2D). Interestingly, SK-N-SH cells overexpressing WT MYCN exhibited a slightly reduced proliferation, accompanied by a morphological change reminiscent of a more epithelial-like phenotype.

Both transient and constitutive overexpression of the MYCN 2D mutant yielded high expression of the product (Figures 2A and 2C), suggesting that PAK2 phosphorylation does not interfere with protein stability. Nevertheless, it could be speculated that the 2D mutation induces an aberrant (e.g. cytosolic) localization of MYCN, therefore not allowing its proper transcriptional activity. To rule out this, SK-N-SH cells expressing the pLV-puro-MYCN WT and 2D constructs were subjected to immuno-fluorescent analysis with an anti-MYCN antibody. As expected, both variants displayed a nuclear localization (Supplementary figure 2). Of note, SK-N-SH cells express c-MYC. As mentioned above, c-MYC and MYCN have been suggested to display a mutually exclusive expression^17^. In this sense, we observed that overexpression of MYCN WT in these cells resulted in a significant downregulation of c-MYC. Interestingly, the MYCN 2D variant did not disrupt c-MYC expression, suggesting that MYCN-dependent downregulation of c-MYC involves MYCN transcriptional activity or repression (Supplementary figure 3). Altogether, our data demonstrates that MYCN and MAX can be phosphorylated by PAK2, and that phosphorylation on 2 conserved PAK2-target sites reduces MYCN:MAX interaction, transcriptional activity, and proliferation of neuroblastoma cells.

**Figure 3.**
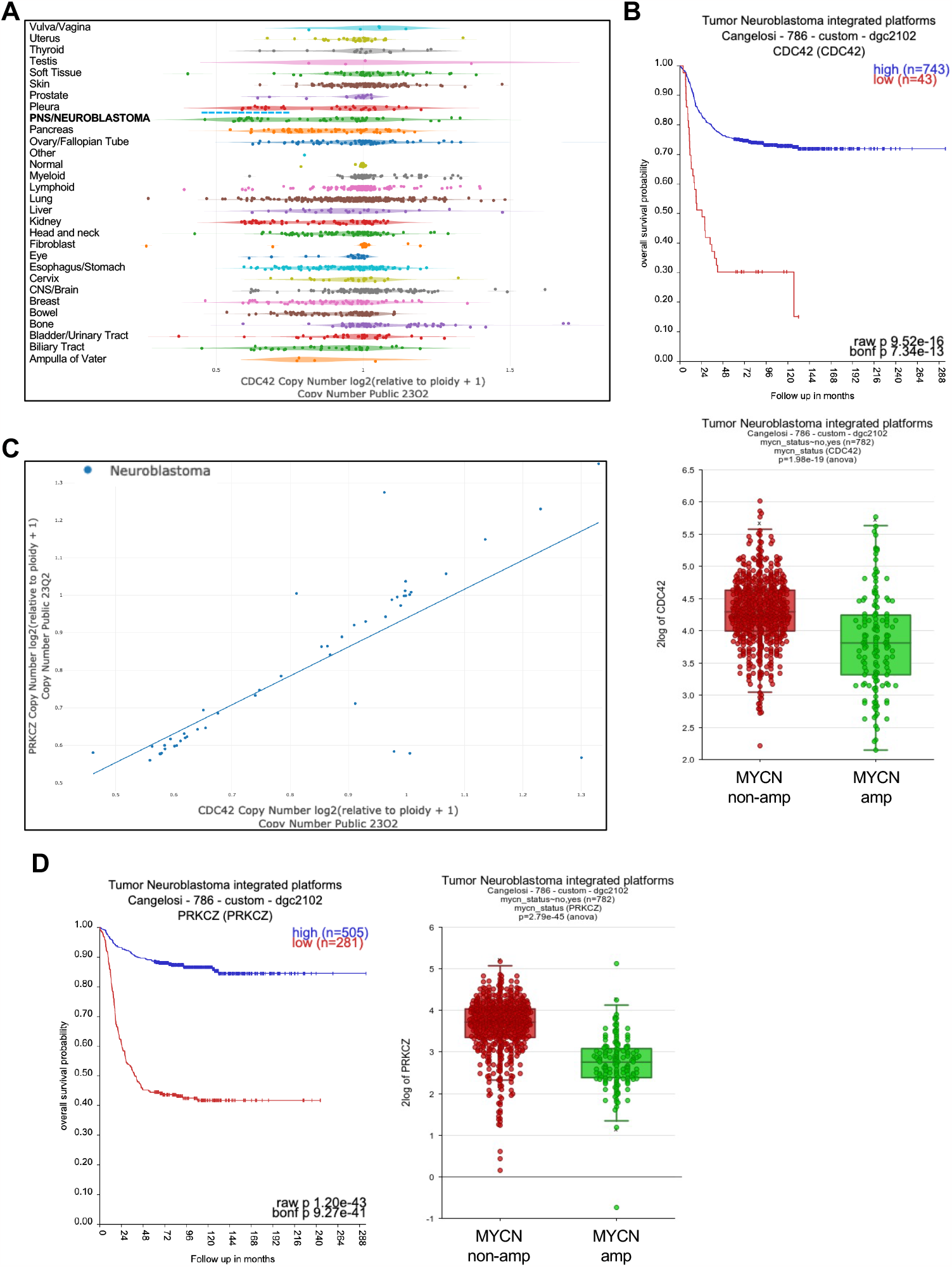
**A**. Analysis of CDC42 copy number across cancer cell lines of the depmap.org portal demonstrates a specially low average copy number in neuroblastoma cell lines, as compared to most other cancer cell lines. **B**. Upper panel. A higher CDC42 expression is significantly correlated to a better neuroblastoma patient survival probability in the Cangelosi 786 cohort from the R2: Genomics Analysis and Visualization Platform. Lower panel. CDC42 expression is significantly lower MYCN-amplified tumors from the Cangelosi 786 cohort (R2: Genomics Analysis and Visualization Platform). **C**. Correlation plot between the copy number of CDC42 and PRKCZ (PKCζ) in Neuroblastoma cell lines from the depmap.org portal. **D**. Upper panel. A higher PRKCZ expression is significantly correlated to a better neuroblastoma patient survival probability in the Cangelosi 786 cohort from the R2: Genomics Analysis and Visualization Platform. Lower panel. PRKCZ expression is significantly lower MYCN-amplified tumors from the Cangelosi 786 cohort (R2: Genomics Analysis and Visualization Platform).

Subsequently, we wondered if PAK2-mediated phosphorylation of MYCN/MAX could be leveraged as a potential strategy to inhibit neuroblastoma cell aggressiveness. PAK2 is a stress-responsive kinase activated by the small GTPases CDC42 and RAC1. Interestingly, CDC42 was previously shown to be downregulated in high-risk neuroblastoma cells and tumors. This was associated to a direct transcriptional inhibition of *CDC42* by MYCN upregulation^36^, but also to the frequent loss of heterozygosity of distal chromosome 1p (1p36del), which is observed in over 70% of MYCN-amplified neuroblastomas^36–38^ and is predictive of an ‘ultra-high-risk’ phenotype^39^. *CDC42* is localized to 1p36.12, and its loss of heterozygosity is also frequently observed in NB cell lines (Figure 3A), Moreover, lower expression of CDC42 is strongly correlated to a worse neuroblastoma patient survival probability (Figure 3B, upper panel), and particularly associated with MYCN-amplified cases (Figure 3B, lower panel). Importantly, restoration of CDC42 expression in MYCN-amplified neuroblastoma cells strongly induced neuronal differentiation^36^, although the mechanisms remain elusive. Therefore, it seems plausible that reduced CDC42 expression in MYCN-amplified neuroblastoma cells results in impaired PAK2 activation and increased MYCN:MAX transcriptional activity. We then wondered if PAK2 could be activated by an alternative mechanism independently of CDC42. As mentioned above, c-MYC Ser373 was also shown to be phosphorylated by PKCζ^32^. Curiously, we noticed that the gene encoding PKCζ -*PRCKZ*-is also localized to 1p36.33 and its copy number in neuroblastoma cell lines is very highly correlated to that of *CDC42* (Figure 3C). Furthermore, lower expression of *PRCKZ* is even more strongly correlated to a worse neuroblastoma patient survival probability than CDC42 (Figure 3D). All these observations suggest that MYCN amplification and 1p36del act cooperatively to repress the tumor-suppressive activity of PAK2, and thus sustain a high MYCN:MAX pro-oncogenic transcription.

We still wanted to explore potential alternative mechanisms for PAK-mediated MYCN:MAX transcriptional inhibition. With this in mind, we searched and annotated the potential correlation of the PAK kinase family members to neuroblastoma patient survival probability. In this sense, we annotated the significant Bonferroni-adjusted p-values for all the expression-to-survival correlations (either positive -associated to better outcomes- or negative -associated to reduced survival-) for all 6 members of the PAK family across 11 neuroblastoma patient cohorts (Figure 4A), as previously described^40^. From this analysis, we could observe that PAK2 did not show a significant association with patient survival, suggesting a minor role in neuroblastoma (Figure 4B, left panel). This is further supported by the fact that PAK2 expression is not particularly enriched in neuroblastoma cell lines as compared to other cancers (Figure 4B, right panel).

**Figure 4.**
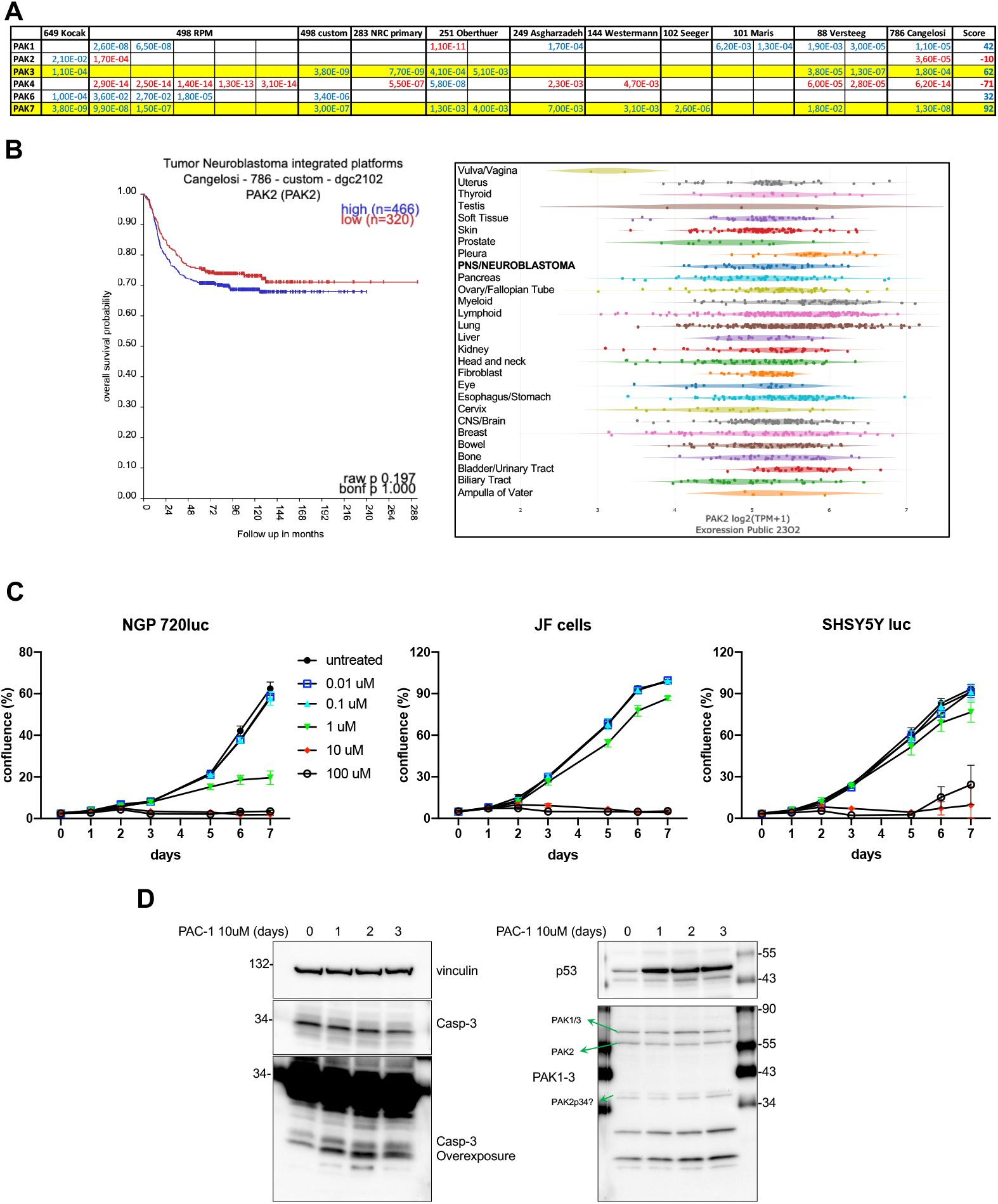
**A**. Bonferroni adjusted p-values (bonf. p-val > 5.00E-02) for the correlations between PAK family members expression and NB patient survival probability across 11 cohorts form the R2: Genomics Analysis and Visualization Platform. Values highlighted in red denote significant negative correlations (higher expression = reduced survival probability), while values highlighted in blue denote positive correlations (higher expression = increased survival probability). Scores are arbitrary units based on the number of datasets showing one or more significant correlations for a given gene, and the value of such significance. We arbitrarily define that a gene must be significantly correlated to either a better or a worse outcome in at least 5 independent cohorts, and obtain a minimum of 50 points (either positive or negative, respectively), to be considered a significant hit. PAK3 and PAK7 show significant positive correlations (highlighted in yellow). **B**. Left panel. Kaplan-Meier plot for the correlation of PAK2 expression and neuroblastoma patient survival probability in the Cangelosi 786 cohort from the R2: Genomics Analysis and Visualization Platform. A higher PAK2 expression is not significantly correlated to a better neuroblastoma patient survival probability. Right panel. Analysis of PAK2 expression across cancer cell lines of the depmap.org portal suggests that PAK2 is not particularly enriched in neuroblastoma cell lines, as compared to most other cancer cell lines. **C**. NGP-luc JF and SHSY5Y-luc neuroblastoma cell lines were plated on M96 well plates (∼2x10^3^ cells/well, 4 replicates/condition) and allowed to grow for 7 days in the absence (untreated) or presence of the indicated final concentrations of the procaspase-3 activating drug PAC-1. Confluency of each well was measured every 1-2 days in an Image cytometer (Celigo, Nexcelom, see Materials and Methods section). **D**. SHSY5Y cells were treated with 10uM PAC-1 for the indicted times and processed for Western blot analysis to detect the indicated proteins.

Interestingly, PAK2 was previously shown to have an alternative activation mechanism independently of CDC42/RAC1, and mediated by direct proteolytic cleavage of PAK2 by activated caspase 3^41,42^. Moreover, small-molecules for the specific activation of caspase 3 have been developed and suggested to have tumor suppressive activities, such as PAC-1^43,44^. In this context, we treated three neuroblastoma cell lines with increasing concentrations of the caspase 3-activating compound PAC-1 and observed a dose dependent inhibition on cell proliferation (Figure 4C), with massive cell death (not shown) at higher concentrations (10-100 μM), probably due to a cytotoxicity effect or to caspase 3-dependent apoptosis, independently of PAK2-mediated MYC(N):MAX inhibition. Although mild effects were seen at 1 μM, these results probably imply a low specific effect in these cells, likely due to a relatively low expression of PAK2. To further clarify this aspect, we performed Western blot analysis of SH-SY-5Y cells treated with 10 μM PAC-1 for up to 3 days. In this context, we detected a mild PAC-1-dependent activation of procaspase 3 (as measured by the appearance of two bands of around 17-19kDa) accompanied by an increase in the levels of p53, but we failed to observe significant changes in the abundance of active PAK2p34, the caspase-3 proteolytic product^42^ (Figure 4D). Noteworthy, the caspase 3 cleavage site in PAK2 is not conserved in other PAK family kinases^41^, suggesting that this alternative mechanism might not be extensive to other PAK family members more relevant to neuroblastoma biology.

Interestingly, PAK3 and PAK7 (a.k.a. PAK5) did show significant positive correlations in 6 and 9 patient datasets, respectively, suggesting that their increased expression might have a tumor suppressive role in neuroblastoma (Figure 4A and Supplementary figure 4), while PAK4 exhibited a strong correlation with a worse outcome in 6 out of 11 neuroblastoma patient cohorts. Furthermore, both PAK3 and PAK7 expression is particularly enriched in neuroblastoma cell lines (Figure 5A and Supplementary figure 5), suggesting that they could be ectopically activated by external stimuli (such as small-molecule compounds) to pharmacologically suppress MYCN:MAX transcription. As a proof-of-concept approach to this idea, we performed a luciferase reporter assay with the pEBOX4-luc system in the presence or absence of WT or 2D MYCN, but also with additional overexpression of PAK3. As expected, expression of pEBOX4-luc/MYCN WT again resulted in maximal luciferase activity, while replacement by MYCN 2D led to a significant reduction of transcriptional response. More importantly, co-expression of PAK3 in the pEBOX4-luc/MYCN WT condition also resulted in a very significant impairment of luciferase expression (Figure 5B), suggesting that PAK3 can directly repress MYCN-dependent transcription, probably by direct phosphorylation of the PAK family conserved sites.

**Figure 5.**
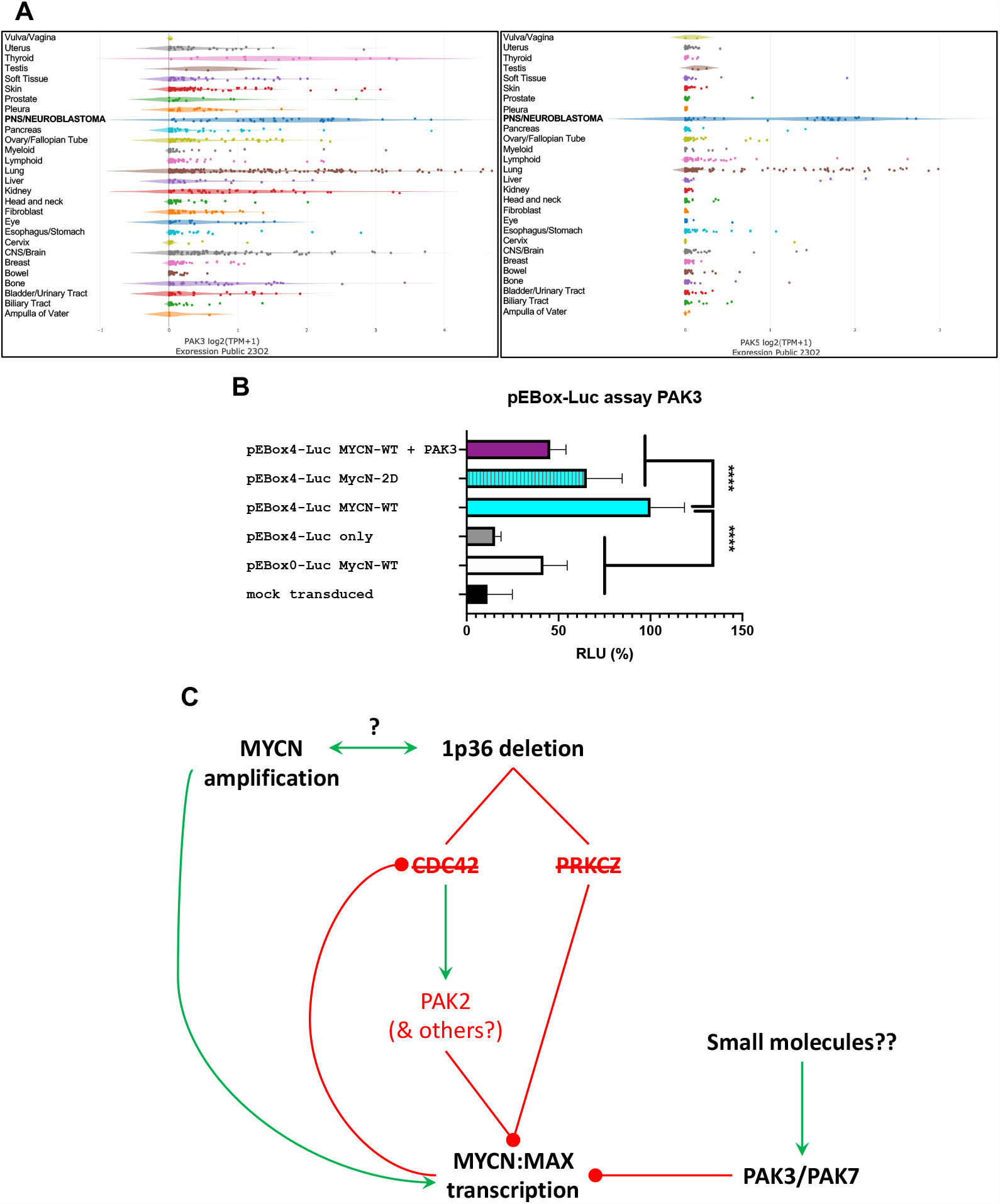
**A**. Analysis of PAK3 (left panel) and PAK5/7 (right panel) expression across cancer cell lines of the depmap.org portal demonstrates that PAK3 and particularly PAK5/7 are highly enriched in neuroblastoma cell lines, as compared to most other cancer cell lines. **B**. Luciferase reporter assay with the pEBOX4-luc system, as described in 2B, in the presence or absence of WT or 2D MYCN, but also with additional overexpression of PAK3. Luciferase activity is expressed in percentual Relative Luciferase Units (RLU %). Each condition was done in 5 technical replicates. **C**. Schematic cartoon of the working model for the regulation of MYC family proteins by the PAK kinases, and its implication for MYCN-driven cancers. MYCN amplification is highly associated with 1p36 deletions although a causal relationship between these two events is unknown. 1p36del results in loss-of-heterozygosity and reduced expression of CDC42 and PRKCZ, two upstream activators of PAK kinases. Therefore, inhibitory phosphorylation of MYCN and MAX by PAK kinases is blunted. Moreover, MYCN can transcriptionally repress CDC42 expression. Finally, PAK3 and PAK7 are highly and exclusively expressed in Neuroblastoma tumors and cells, and PAK3 is shown to repress MYCN transcriptional activity. Therefore, we propose that pharmacological activation of PAK3/PAK7 might constitute a novel therapeutic strategy for the treatment of c-MYC and MYCN-mediated neuroblastoma and associated aggressive cancers.

## Discussion

Here we present robust evidence supporting the existence of a conserved endogenous mechanism for the negative regulation of the MYC family of transcription factors by the CDC42/PAK kinases module, with a special focus on the modulation of MYCN in high-risk neuroblastoma. First, we show that, like c-MYC, MYCN and -importantly-MAX can also be phosphorylated by PAK2 in 2 conserved residues, resulting in the disruption of MYCN:MAX interaction and transcriptional activity. We further show that these phosphorylation events could lead to a complete impairment of neuroblastoma cell proliferation, suggesting a potential new therapeutic avenue for the treatment of MYCN-dependent neuroendocrine and pediatric tumors. We also observed that the activation of the PAK kinases in high-risk neuroblastoma cells is likely suppressed by the cooperative effect of MYCN overexpression and 1p36 deletion, by means of reduced expression of the upstream PAK activators CDC42 and PKCζ (Figure 5C). Finally, we provide important data suggesting that the PAK family members PAK3 and PAK7 are highly expressed in neuroblastoma cells and that at least PAK3 also holds the ability to suppress MYCN transcriptional activity, opening a novel mechanistic avenue for the potential inhibition of MYCN-dependent high-risk neuroblastoma through pharmacological activation of PAK3/PAK7 (Figure 5C). PAK3 and PAK7 are two largely understudied kinases, and very little is known about their physiological mechanisms of activation and consequences. Even less is known about potential alternative or artificial mechanisms for activation of these kinases. Therefore, future efforts should focus on better understanding the molecular pathways governing PAK3 and PAK7 and on how to leverage new pharmacological tools for the direct activation of these kinases as an innovative approach for the potential treatment of MYCN-amplified neuroblastoma and other associated aggressive neuroendocrine tumors.

## Materials and Methods

### Cell lines and culture

All cell lines were cultivated at 37 °C with 5% CO_2_. HEK-293T cells were maintained in Dulbecco’s modified Eagle medium (DMEM), supplemented with 10% FCS and antibiotics (100 units/ml penicillin and 100 μg/ml streptomycin). The NGP cell line was obtained from DSMZ and cells were grown in DMEM (4.5 g/L glucose) + 10% h.i. FBS + 4mM L-Glutamine + pen/strept. according to vendors’ recommendations and passaged no more than 8 times. The neuroblastoma cell line SJNB-JF-G12 (JF) was originally established in 1979 from a patient with disseminated neuroblastoma and was a kind gift from Dr. Malcom Brenner. JF cells were grown in RPMI 1640 + 10% h.i. FBS + 4mM L-Glutamine + pen/strept. SH-SY5Y cells were obtained from ATCC, and were grown in DMEM/F12 (1:1) + 10% h.i. FBS + 4mM L-Glutamine + penicillin/streptomycin. All cell lines were tested for Mycoplasma every 2 months using e-Myco™ Mycoplasma PCR Detection Kit (Bulldog Bio #25235), according to the manufacturer’s instructions. All cell lines were authenticated by STR analysis (ATCC).

### Reagents

Procaspase activating compound 1 (PAC-1) was obtained from MedChemExpress (HY-13523) and resuspended to a 10mM stock in DMSO for *in vitro*/cell culture-based assays. Primary antibodies from Cell Signaling Technology were: FLAG-tag (#14793), HA-tag (#3724), vinculin (#13901), MYCN (#84406), c-MYC (#18583), PAK1/2/3 antibody (#2604) and Caspase-3 Antibody (#9662). Other primary antibodies were: Monoclonal anti-FLAG^®^ M2 from Millipore Sigma (F1804), c-MYC (sc-40) and MAX (sc-8011) from Santa Cruz Biotechnology. Mouse anti-rabbit (211-035-109) and goat anti-mouse (115-035-146) HRP-conjugated secondary antibodies were from Jackson ImmunoResearch Laboratories Inc. Constitutively activated mutant recombinant PAK2 (T402E) plasmid was a kind gift from Dr. Jeanne Hardy. The recombinant protein was bacterially expressed and purified following the protocol described in ^45^. Dynabeads Protein A/G-magnetic beads for immuno-precipitation were obtained from Invitrogen (#10001D/10003D). Lipofectamine 2000 transfection reagent was from Invitrogen (#11668027). pCDNA3-HA-human MYCN expression construct was obtained form Addgene (#74163). pFLAG-MAX WT (p21) expression construct was a kind gift from Dr. Julia Weigand^46^. The pRK5-myc-PAK3 was a kind gift from Dr. Richard Huganir^47^. Respective 2D mutant versions of HA-MYCN and FLAG-MAX were obtained by standard molecular biology techniques using the QuickChange mutagenesis kit as recommended by the manufacturer. pLV-puro-MYCN WT and 2D constructs for lentiviral expression were obtained by direct cloning of the pCDNA3-HA-MYCN constructs into the pLV-puro vector by standard molecular biology techniques. All constructs were confirmed by Sanger sequencing. The MYCN:MAX luciferase reporter plasmids pEBOX0 (pMyc0ElbLuc #53242) and pEBOX4 (pMyc4ElbLuc #53246) were obtained from Addgene, as well as pRenilla (#118059). Luciferase reporter assays were performed with Dual-Luciferase^®^ Reporter Assay System from Promega as recommended by the manufacturer.

### Western blot

Western blot analysis was conducted using standard methods^48^. Briefly, cells grown to a 60-80% confluency were lysed in radio-immuno-precipitation assay (RIPA) lysis buffer (Prometheus Protein Biology Products #18-416) or IP Buffer (see below) supplemented with Protease and Phosphatase Inhibitor Cocktails (Pr/Ph-ICs; Pierce, Thermo Scientific A32955 and A32957). Lysates were sonicated on ice, centrifuged at 15,000×g at 4 °C for 20 minutes and the soluble protein fraction was collected. Protein extracts were quantified using a Pierce BCA Protein Assay Kit (Thermo Scientific #23227). A total of 30-50 μg of proteins were separated via SDS-PAGE using Novex™ WedgeWell™ 4-20%, Tris-Glycine Mini Protein Gels (Invitrogen, Thermo Scientific XP04202BOX) and blotted onto a PVDF membrane using an iBlot transfer system and transfer stacks (Invitrogen, Thermo Scientific IB401001). Proteins were detected using SuperSignal™ West Pico PLUS Chemiluminescent Substrate (Thermo Scientific 34580). A ChemiDoc MP Imaging System (Bio-Rad) was used for chemiluminescent detection and analysis.

### Immuno-precipitations

For soluble cell lysates, cells were washed twice in PBS and lysed in IP buffer (50 mM Tris-HCl [pH 7.4], 150 mM NaCl, 2 mM EDTA, 1% NP-40, 10% glycerol, Pr/Ph-ICs). Lysates were clarified by centrifugation and protein quantification with the BCA assay kit (Pierce). One mg of total protein/sample were incubated for 6 h at 4°C with protein A/G-magnetic beads (Dynabeads, Invitrogen) prebound with 5 μg of the corresponding antibody, and then beads were washed with IP buffer. Finally, both lysates (input) and the immuno-precipitates were resuspended in 5X LB and analyzed by Western blotting.

### *In vitro* kinase assays

For *in vitro kinase* (IVK) from immuno-complexes, cell lysates were prepared in IP lysis buffer (50 mM HEPES [pH 7.4], 75 mM NaCl, 1 mM EDTA, 1% NP-40, Pr/Ph-ICs). Cell lysates were incubated overnight at 4°C with the corresponding antibody bound to protein A/G-magnetic beads (Dynabeads, Invitrogen). Immuno-complexes were washed 4 times with IP and twice time with kinase buffer (25 mM Hepes pH 7.4, 5 mM MgCl2, 5 mM MnCl2, 0.5 mM DTT). Immuno-complexes were split into 3 aliquots: 5-10% for Western blotting and 2 aliquots of ∼45% for IVK assay, with and without human recombinant PAK2. Immuno-complexes were incubated for 20 min at 30°C in 30 μl of kinase buffer with a final concentration of 50 μM ATP and [γ-^32^P] ATP (1 x 10^−2^ μCi/pmol; PerkinElmer). Reactions were stopped by adding 5X LB, and samples were resolved by SDS-PAGE and then stained with Coomassie blue. ^32^P incorporation was detected by autoradiography of dried gels.

### Cell proliferation analysis

Cell proliferation of indicated conditions (untreated vs. MYCN WT/2D-expressing cells) was measured as the relative whole-well confluency of 96-well culture plates using a Celigo Imaging Cytometer (Nexcelom Bioscience LLC). Briefly, ∼2000 cells/well were plated on day -1 and incubated for 24h to allow the cells to attach and recover. Relative confluence was subsequently analyzed every 24/48hs until untreated control wells reached 70-90% confluency. Each condition was done in 4-6 replicates per experiment. Medium was replaced every 48-72h. Cell proliferation was plotted as the time-dependent change of the average relative confluency for each condition using GraphPad Prism 8 software.

### Immuno-fluorescence analysis

Standard immuno-fluorescence techniques were used as recommended in the Cell Signaling Technology protocols webpage. Briefly, cells were fixed in 4% paraformaldehyde in PBS for 15 min at room temperature, washed 2X in PBS (5 min/wash) and 2X with 0.2% Triton X-100 in PBS (‘PBS-T’). Cells were blocked in 2% BSA-PBS-T (blocking buffer) for 60 min at 4ºC and incubated with primary antibodies diluted in blocking buffer overnight at 4ºC. Then, 4X washes in PBS and 1h incubation with secondary antibody Alexa 488 goat anti-rabbit (Invitrogen) followed by 4X additional washes in PBS. Images were obtained in an Echo Revolve fluorescence microscope (Bico) or in a Zeiss LSM700 confocal microscope (UMass Chan Medical School).

### Lentivirus preparation and infection

HEK-293T cells were transfected with pVSV-G^49^ and pCMVΔR8.91^50^, together with the pLV-puro-MYCN WT/2D vectors using Lipofectamine™ 2000 reagent (Invitrogen) as recommended by the fabricant, and following the recommendations of The RNAi Consortium (TRC) laboratory protocols with slight modifications. Twelve hours after transfection the medium was replaced by DMEM, supplemented with 30% FCS and antibiotics which. Cell supernatants were harvested every 24 hs, replacing with fresh medium and stored at 4ºC until collection of the last harvest (at 72 hs). At this point, the consecutive harvests were pooled, filtered through 0.45 μm filters and split in 3-5 mL aliquots, which were stored at -80ºC. NB cells were infected with shControl or shRNA lentiviral particles by adding a 1:1 mix of medium:viral supernatant for 24-48 hs. Puromycin selection (2 μg/ml) was applied for 2-3 days and always compared to non-transduced control cells, which generally died within the first 24 hs. Target overexpression was confirmed by Western Blot.

### Statistical Analyses

All quantitative data points represent the mean of 2-3 independent experiments performed in 3 or more replicates with standard deviation (S.D). Statistical analysis was performed using t-test or two-way ANOVA (GraphPad Software, Inc., La Jolla, CA).

