## Supplementary figures 1-5 for "Phosphorylation of MYCN and MAX by PAK family kinases is a novel tumor-suppressor mechanism in neuroblastoma with potential therapeutic implications"

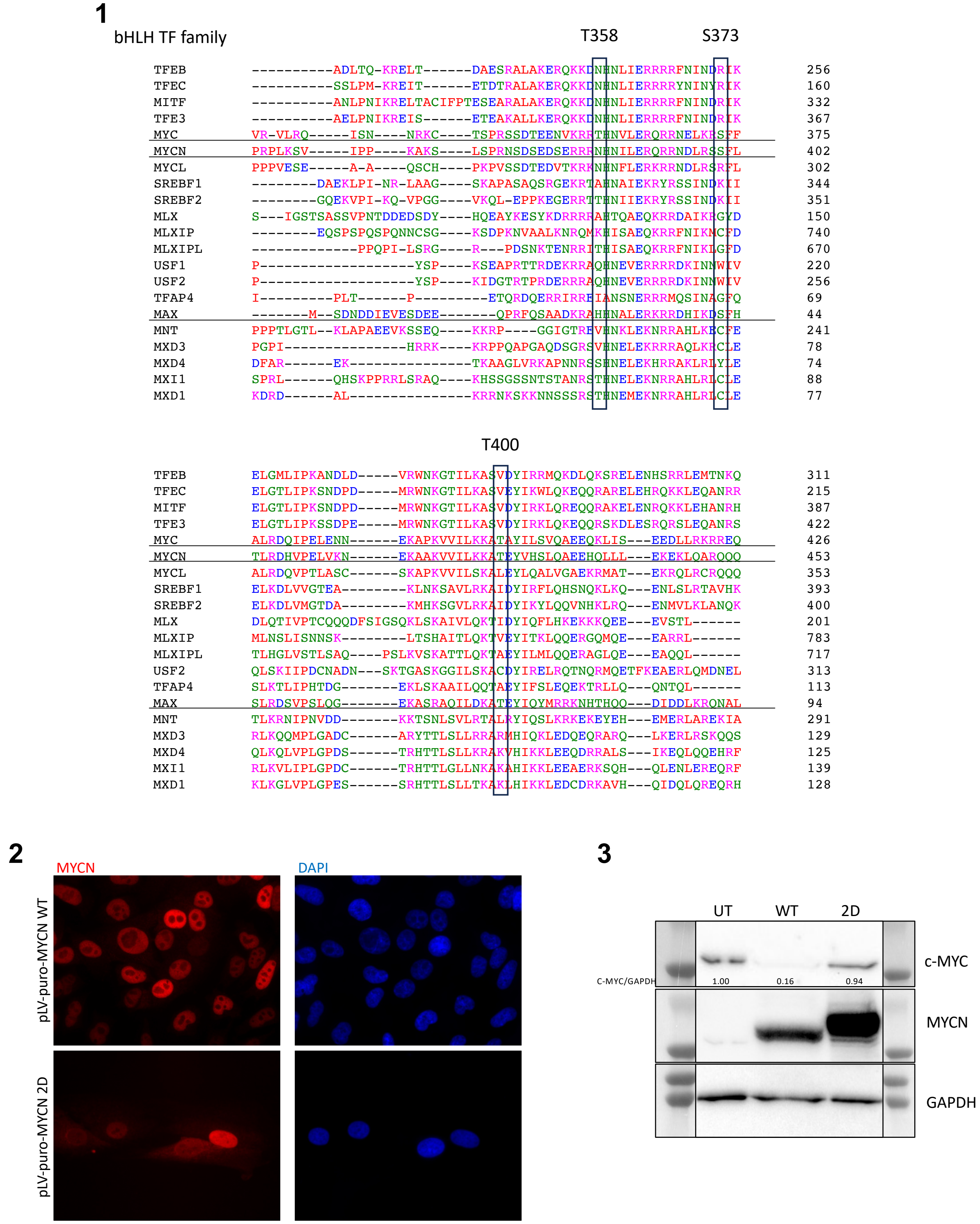

4

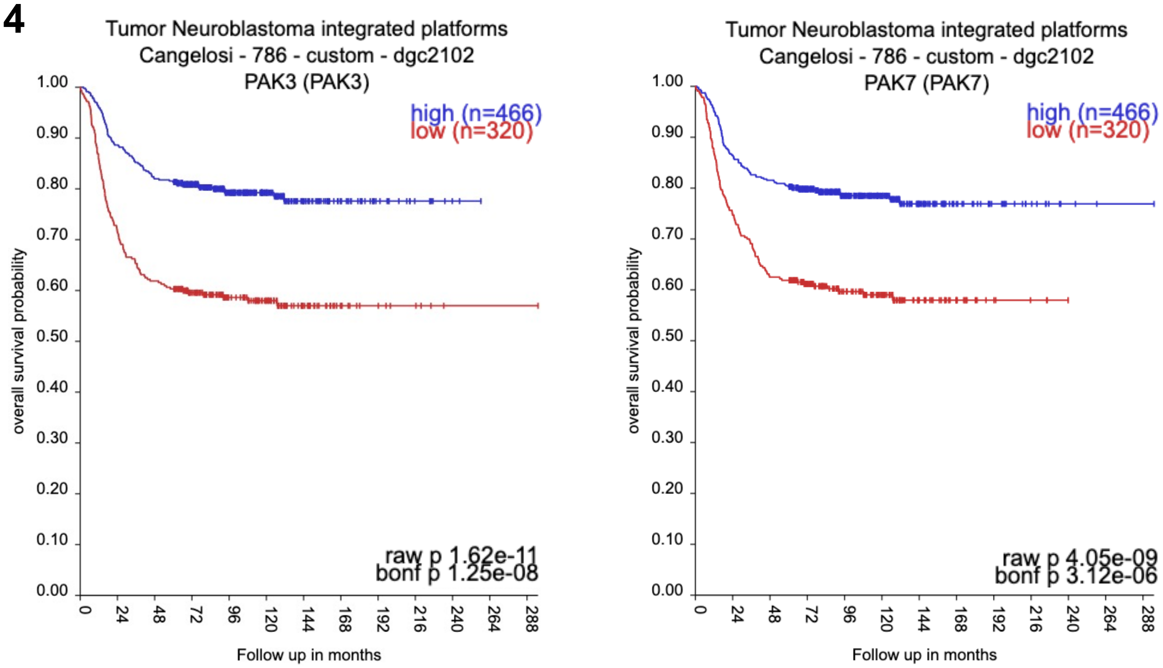


5

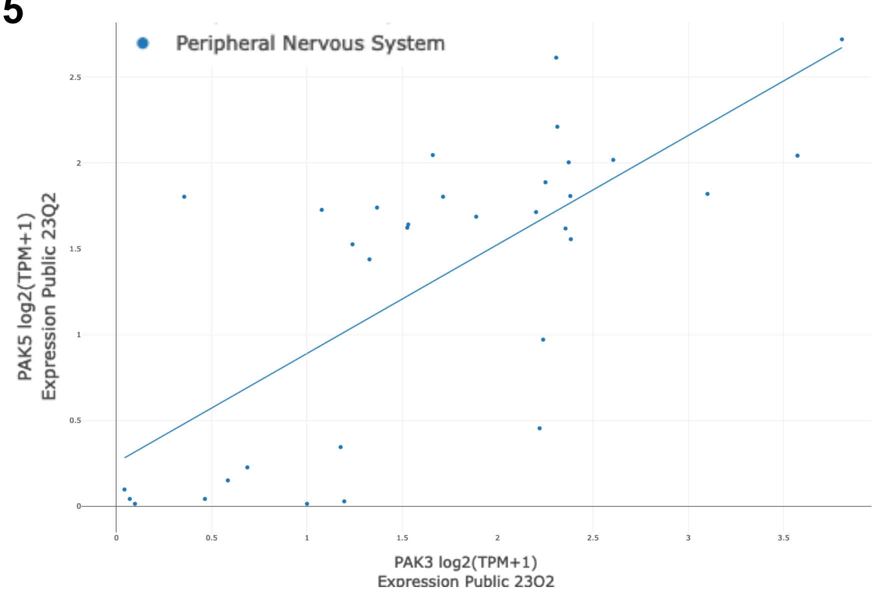


### Supplementary figure legends

**1.** Related to Figure 1B. Partial amino-acid sequence alignment of human bHLH/Z family members depicting the conservation of PAK2 target sites (black boxes) at the level of c-MYC Thr358, Ser373 and Thr400.

**2.** Related to Figure 2 C-D. SK-N-SH neuroblastoma cells were transduced with pLV-puro-MYCN WT or 2D lentiviruses. Upon puromycin selection, cells were processed for immunostaining with anti-MYCN antibody, demonstrating a nuclear MYCN localization in both conditions.

**3.** Related to Figure 2 C-D. SK-N-SH neuroblastoma cells were transduced with pLV-puro-MYCN WT or pLV-puro-HA-MYCN 2D* lentiviruses. Upon puromycin selection, cells were processed for Western blot analysis for the indicated proteins (c-MYC, MYCN and GAPDH). * MYCN band in the 2D condition displays a shift due to the use of a HA-tagged overexpression variant in this experiment, while the WT version did not have a tag.

**4.** Related to Figure 4A. Kaplan-Meier plots for the correlation of PAK3 and PAK5/7 expression and neuroblastoma patient survival probability in the Cangelosi 786 cohort from the R2: Genomics Analysis and Visualization Platform. Higher PAK3/7 expression is significantly correlated to a better neuroblastoma patient survival probability.

**5.** Related to Figure 5 A. Correlation plot between the expression of PAK3 and PAK5/7 in neuroblastoma cell lines from the depmap.org portal.
